# Evolution of the DAN gene family in vertebrates

**DOI:** 10.1101/794404

**Authors:** Juan C. Opazo, Federico G. Hoffmann, Kattina Zavala, Scott V. Edwards

## Abstract

The DAN gene family (DAN, Differential screening-selected gene Aberrant in Neuroblastoma) is a group of genes that is expressed during development and plays fundamental roles in limb bud formation and digitation, kidney formation and morphogenesis and left-right axis specification. During adulthood the expression of these genes are associated with diseases, including cancer. Although most of the attention to this group of genes has been dedicated to understanding its role in physiology and development, its evolutionary history remains poorly understood. Thus, the goal of this study is to investigate the evolutionary history of the DAN gene family in vertebrates, with the objective of complementing the already abundant physiological information with an evolutionary context. Our results recovered the monophyly of all DAN gene family members and divide them into five main groups. In addition to the well-known DAN genes, our phylogenetic results revealed the presence of two new DAN gene lineages; one is only retained in cephalochordates, whereas the other one (GREM3) was only identified in cartilaginous fish, holostean fish, and coelacanth. According to the phyletic distribution of the genes, the ancestor of gnathostomes possessed a repertoire of eight DAN genes, and during the radiation of the group GREM1, GREM2, SOST, SOSTDC1, and NBL1 were retained in all major groups, whereas, GREM3, CER1, and DAND5 were differentially lost.

## Introduction

Genetic variation harbored in non-model species represents a powerful resource to gain insights in our understanding of the genetic bases of biological diversity (Opazo et al. 2005; Storz et al. 2007; Faulkes et al., 2015). The comparative approach has gained popularity since the discovery of non-model species that are resistant to diseases that affect human health, and also because some non-model species develop pathologies in a similar way as ourselves (Protas et al. 2006; Gorbunova et al., 2012; Castro-Fuentes and Socas-Pérez, 2013; Manov et al., 2013; Henning et al., 2014; Braidy et al., 2015).

The DAN gene family (DAN stands for Differential screening-selected gene Aberrant in Neuroblastoma) is a group of genes characterized by the presence of a DAN domain (http://pfam.xfam.org/family/PF03045), low inter-paralog conservation and expression during early development. DAN genes play fundamental roles in limb bud formation and digitation, kidney formation, and morphogenesis as well as left-right axis specification (Nolan & Thompson, 2014). In adult organisms, the expression of DAN genes is associated with pathologies such as cancer and nephropathies (Koli et al. 2006; Walsh et al. 2010; Gu et al. 2012; Droguett et al. 2014; Liang et al. 2015). As currently recognized, the DAN gene family comprises seven paralogs: NBL1 (neuroblastoma 1); CER1 (cerberus 1); DAND5 (DAN domain BMP antagonist family member 5); SOST (sclerostin); SOSTDC1 (sclerostin domain containing 1); GREM1 (gremlin 1); and GREM2 (gremlin2) (Avsian-Kretchmer et al. 2004; Walsh et al. 2010; Nolan & Thompson, 2014). Historically, DAN genes have been associated with the inhibition of the bone morphogenetic protein (BMP) signaling pathway, although more recent studies have shown that they also act as antagonists to nodal, wingless-type MMTV integration site (Wnt) and vascular endothelial growth factor (VEGF) signaling cascades (Piccolo et al. 1999; Ellies et al. 2006; Chiodelli et al. 2011). Most of the work on DAN genes has focused on understanding their role in development (Nolan & Thompson, 2014, and references therein), and less effort has focused on their evolutionary history (Avsian-Kretchmer et al. 2004; Walsh et al; 2010; Nolan & Thompson, 2014; Le Petillon et al. 2013; Opazo et al. 2017). Understanding the evolutionary history of gene families permits insights into the ways in which genes originate and diversify, as well as rates of evolution, the relative roles of different mutational processes such as point mutations and duplications and, most importantly, relating all of these phenomena to the physiological and life-history idiosyncrasies of different taxonomic groups.

The goal of this study is to investigate the evolutionary history of the DAN gene family in vertebrates, with the aim of complementing the already abundant physiological information with an evolutionary perspective. Therefore, we examined the diversity of DAN genes in representative species of all major groups of vertebrates, inferred homologous relationships by examining gene phylogenies and synteny conservation, reconstructed ancestral gene repertoires, examined patterns of differential gene retention and quantified rates of molecular evolution. Our results recovered the monophyly of all DAN gene family members and grouped them into five clades. In addition to the well-known DAN genes, our phylogenetic results revealed the presence of two additional DAN gene lineages; one is only retained in cephalochordates (e.g. amphioxus), whereas the second one (GREM3) has only been retained by cartilaginous fish, holostean fish, and coelacanth. The phylogenetic distribution of genes suggests that the ancestor of gnathostomes possessed a repertoire of eight DAN genes, and that during the radiation of vertebrates, GREM1, GREM2, SOST, SOSTDC1, and NBL1 were retained in all major groups, whereas, GREM3, CER1, and DAND5 were differentially lost, creating complex patterns of retention and diversification.

## Methods

### DNA data and phylogenetic analyses

*To minimize potential errors in the annotation of the sequences, w*e manually annotated DAN genes in representative species of all major groups of chordates. To do so, we first identified genomic fragments containing DAN genes in Ensembl v86 (Zerbino et al., 2018) or National Center for Biotechnology Information (NCBI) databases (refseq_genomes, htgs, and wgs; Geer et al., 2010) based on conserved synteny s. Once identified, genomic pieces were extracted including the 5’ and 3’ flanking genes. After extraction, we manually curated the existing annotation or we annotate the novo by comparing known exon sequences (query sequence) from a species that share a common ancestor most recently in time to the species of which the genomic piece (subject sequence) is being analyzed using the program Blast2seq v2.5 with default parameters (Tatusova & Madden, 1999). Sequences derived from shorter records based on genomic DNA or complementary DNA were also included to attain a broad taxonomic coverage. We included representative species from mammals, birds, reptiles, amphibians, lobe-finned fish, holostean fish, teleost fish, cartilaginous fish, cyclostomes, urochordates and cephalochordates (Supplementary Table S1). Amino acid sequences were aligned using the FFT-NS-i strategy from MAFFT v.6 (Katoh & Standley, 2013). Our alignment contained 424 sequences and a length of 797 characters. Phylogenetic relationships were estimated using maximum likelihood and Bayesian approaches. Our goal was to estimate the history of a gene family, not the phylogeny of the organismal lineages themselves, hence we used traditional gene-tree based phylogenetic methods (Edwards, 2009; Yang & Rannala, 2012). We used IQ-Tree (Kalyaanamoorthy et al., 2017) to select the best- fitting model of amino acid substitution (JTT + I + G4) and to obtain the maximum likelihood tree (Trifinopoulos et al. 2016) and assessed support for the nodes with 1,000 bootstrap pseudoreplicates using the ultrafast routine. Bayesian searches were conducted in MrBayes v.3.1.2 (Ronquist and Huelsenbeck, 2003), setting two independent runs of six simultaneous chains for 5×10^6^ generations, sampling every 2,500 generations, and using default priors. Once the analyses were done, we verified that the estimated sample size (ESS) exceeded the recommended value of 200 using Tracer ver 1.7.1 (Rambaut et al. 2018). The run was deemed converged once the likelihood scores reached an asymptotic value and the average standard deviation of split frequencies remained < 0.01. We discarded all trees that were sampled before convergence, and we evaluated support for the nodes and parameter estimates from a majority rule consensus of the last 4,000 trees.

### Assessments of Conserved Synteny

We examined genes found upstream and downstream of each member of the DAN gene family on species representative of all the major groups of vertebrates. Synteny analyses were performed in humans, chicken, spotted gar and elephant shark. Initial ortholog predictions were derived from the EnsemblCompara database (Vilella et al. 2009) and were visualized using the program Genomicus v84.01 (Muffato et al. 2010). In the case of the elephant shark the genomic segments containing DAN genes were annotated, and predicted genes were then compared with the non-redundant protein database using Basic Local Alignment Search Tool (BLAST) (Altschul et al. 1990).

## Results and discussion

### Gene phylogenies and synteny analyses define homology

Our phylogenetic analyses recovered the monophyly of each members of the genes in the DAN gene family with strong support with the exception of DAND5 (Fig. 1). In addition, our trees suggest an arrangement in which these genes are divided into five main groups (Fig. 1): 1) a clade containing the SOSTDC1 and SOST genes; 2) a clade that contains the CER1 and DAND5 gene lineages; 3) a clade corresponding to 2 cerberus-like sequences of cephalochordates; 4) a clade corresponding to the NBL1 gene; and 5) a clade containing the GREM gene lineages (Fig. 1).

**Figure 1.**
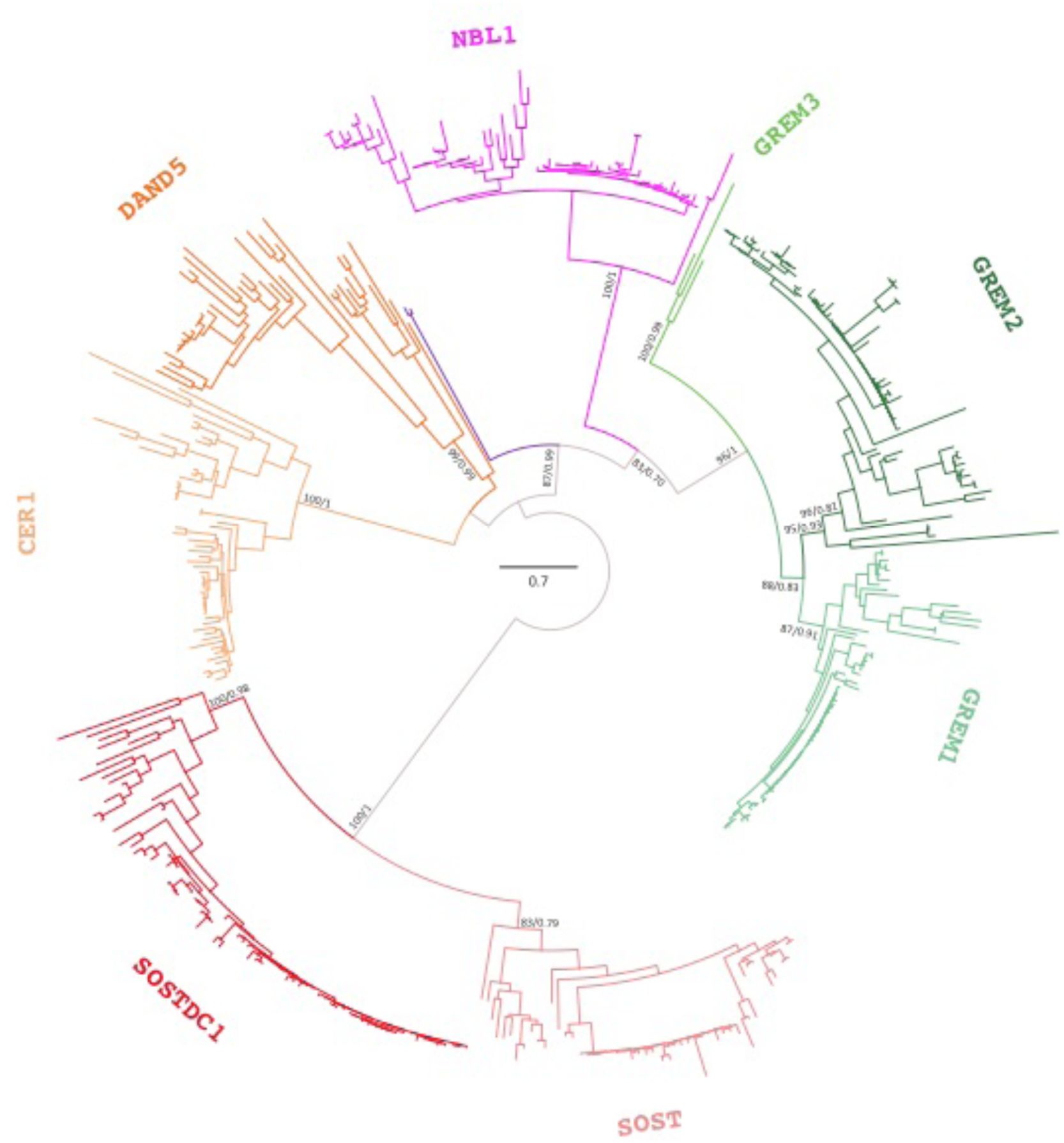
Midrooted maximum likelihood phylogenetic tree depicting relationships among DAN genes. Numbers on the nodes correspond to maximum likelihood ultrafast bootstrap support and Bayesian posterior probability values. The bar represents the number of amino acid substitutions per site.

In the first clade, the monophyly of SOSTDC1 and SOST is well supported, as is the sister group relationship between them (shown in shades of red; Fig. 1), suggesting that they share a common ancestor more recently than with any other member of the family (Supplementary Fig. S1A). The sister group relationship between these two gene lineages has also been recovered in other studies (Avsian-Kretchmer et al. 2004; Walsh et al. 2010; Nolan & Thompson, 2014). Synteny analyses provide further support for the identity of these two gene lineages because genes found up- and downstream are conserved in most surveyed species (Fig. 2). The tree topology in which the SOSTDC1/SOST clade is recovered sister to all other members of the gene family has also been recovered in other studies (Avsian-Kretchmer et al. 2004; Walsh et al. 2010; Nolan & Thompson, 2014).

**Figure 2.**
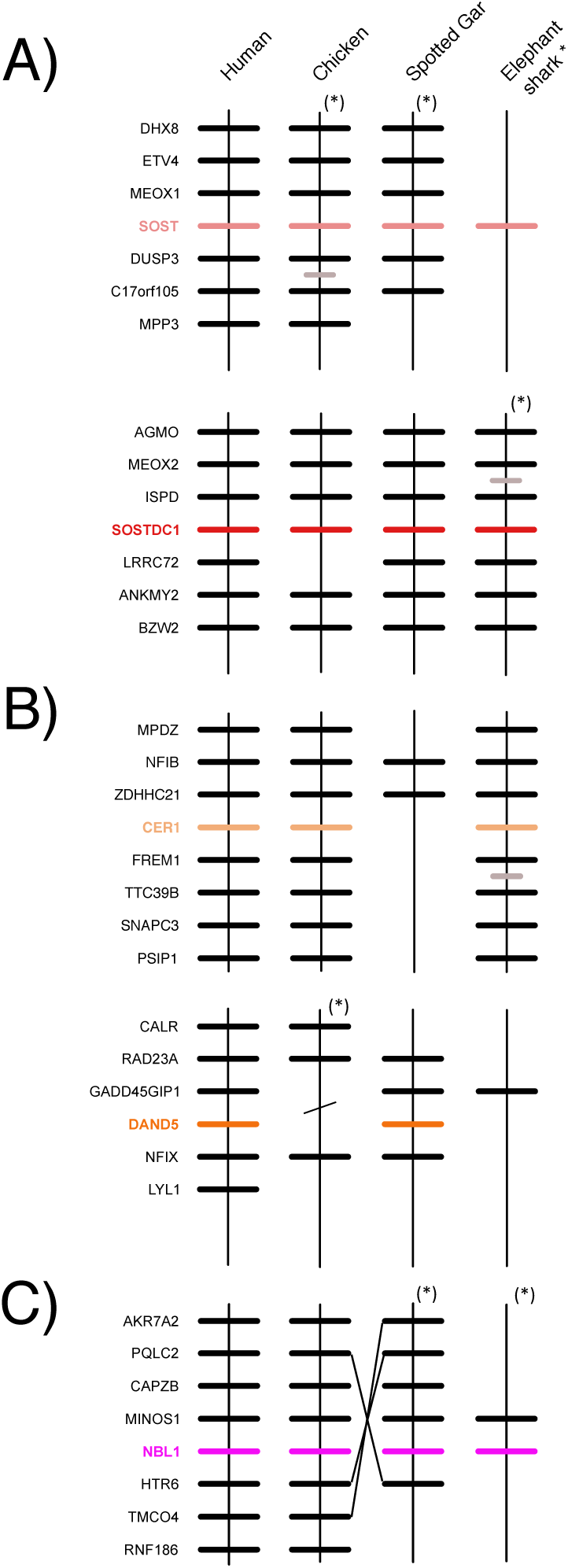
Patterns of conserved synteny in the chromosomal regions harboring DAN genes. A) Chromosomal region harboring SOST and SOSTDC1 genes; B) region harboring the CER1 and DAND5 genes; C) Chromosomal region harboring the NBL1 genes. Asterisks denote that the orientation of the genomic piece is from 3′ to 5′, gray lines represent intervening genes that do not contribute to conserved synteny, diagonals denote that the chromosomal pieces are not continuous, whereas the lack of a line denotes the absence of the gene.

In the second group, shown in shades of orange (Fig.1), the monophyly of CER1 genes is well supported, whereas the monophyly of DAND5 genes is not (Fig. 1). The lack of support for the DAND5 clade could be due to the presence of a coelacanth DAND5 sequence for which the phylogenetic position is not well resolved (Supplementary Fig. S1B). However, genes found upstream (GADD45GIP1 and RAD23A) and downstream (NFIX, LYL1) of the coelacanth DAND5 sequence confirm that the gene is in the expected position for a DAND5 ortholog, providing independent support for the topology obtained in our phylogenetic analyses (Fig. 1). Thus, the data suggests that the coelacanth DAND5 gene is a true ortholog of the DAND5 gene of vertebrates and that the lack of resolution is likely due to high sequence divergence in the coelacanth gene. In addition to the coelacanth sequence, synteny analyses provide further support for our phylogeny, because genes found up- and downstream of CER1 and DAND5 are conserved across species (Fig. 2). Although we recovered a sister group relationship between CER1 and DAND5 (Fig. 1), the relevant node was not supported (Fig. 1). In the literature, there is no clear pattern regarding the sister group relationship between these paralogs (Avsian-Kretchmer et al. 2004; Walsh et al. 2010; Nolan & Thompson, 2014). Although the study of Walsh et al. (2010) supports the sister group relationship between CER1 and DAND5, the study of Avsian-Kretchmer et al. (2004) recovered a topology in which CER1 is the sister group of a clade containing DAND5, NBL1 and GREM gene lineages, whereas, Nolan & Thompson (2014) recovered DAND5 sister to a clade containing CER1 and GREM gene lineages. In support of our topology, an amino acid alignment that includes all human DAN family members shows that CER1 and DAND5 are more similar to each other than to any other member of the gene family; moreover, the Ensembl and gene cards platforms suggest that the only paralog of CER1 is DAND5 and vice versa. Finally, according to our analyses, the clade that includes CER1 and DAND5 was recovered sister to the clade that includes the NBL1 and GREM gene lineages and a third clade of DAN genes that includes two cerberus-like sequences from two lancelet species (Fig. 1).

Our phylogenetic analyses recovered a third clade of DAN genes that includes two cerberus-like sequences from two lancelet species (*Branchiostoma floridae* and *B. belcheri*; purple lineage; Fig. 1). These sequences were recovered sister to the GREM/NBL1 clade (Fig. 1). This phylogenetic arrangement would be compatible with two possible evolutionary scenarios. The first suggests that the DAN gene repertoire of lancelets represents the gene complement of the ancestor of Olfactores, from latin olfactus, the group that includes vertebrates and urochordates, and during the radiation of the vertebrates the ancestral gene lineage gave rise to the extant GREM and NBL1 genes. The second scenario implies that the lancelet gene lineage could be an ancient member of the DAN gene family that has only been retained in cephalochordates. In support of the second scenario, we found cephalochordate sequences in the NBL1 and GREM clades (Supplementary Fig. S1C), suggesting that the genomes of chordates possess an already-differentiated member of the GREM and NBL1 clades.

The fourth clade corresponds to the NBL1 gene which was recovered with strong support (pink clade; Fig. 1)(Supplementary Fig. S1C). The conservation of the genes found up- and downstream provides further support for the identity of the NBL1 gene lineage (Fig. 2). The phylogenetic position of NBL1 is still a matter of debate because different phylogenetic hypotheses have been proposed in earlier studies. Nolan et al. (2014) recovered NBL1 as sister to the clade containing GREM1, GREM2, CER, and DAND5 gene lineages; whereas in Walsh et al. (2010) NBL1 sequences were recovered in a trichotomy with the GREM1/GREM2 and CER1/DAND5 clades. However, in support of our study Avsian-Kretchmer et al. (2004) recovered NBL1 as sister to the GREM gene lineages. The fifth clade corresponds to a well-supported group containing GREM gene lineages (shown in shades of green; Fig. 1). Within this clade, the monophyly of the GREM1 and GREM2 genes is well supported (Fig. 1). As in other cases, synteny analyses support the identity of both gene lineages (Fig. 3). We identified a clade that contains a single copy gene in lancelets and tunicates, which in turn was recovered as sister to the GREM2 clade (Supplementary Fig. S1D). The presence of a single copy gene in cephalochordates and urochordates is expected given that a previous study reported that the vertebrate GREM genes diversified as a product of the two rounds of whole-genome duplications early in vertebrate evolution (Singh et al. 2015; Sacerdot et al. 2018; Simakov et al. 2020), and cephalochordates and urochordates diverged from vertebrates before these whole-genome duplications. Unexpectedly, our phylogenetic analyses identified the presence of a third GREM gene lineage (GREM3) among gnathostome vertebrates that has not been described before (Fig. 1). This new gene lineage was recovered with strong support, and was identified in three distantly related species: elephant shark (*Callorhinchus milii*), spotted gar (*Lepisosteus oculatus*) and coelacanth (*Latimeria chalumnae*). Synteny analyses provide further support for the identity of this new lineage as genes found up- (RASGRP4, FAM98C and SPRED3) and downstream (RYR1, MAP4K1, and EIF3K) are conserved (Fig. 3). Moreover, we found that human orthologs of the genes syntenic to GREM3 mapped to chromosome 19, suggesting that this chromosome as the putative genomic location of the GREM3 gene in humans (Fig. 3). A similar differential retention of an ancestral gene in a small group of distantly related species has also been observed in other gene families (Wichmann et al. 2016).

**Figure 3.**
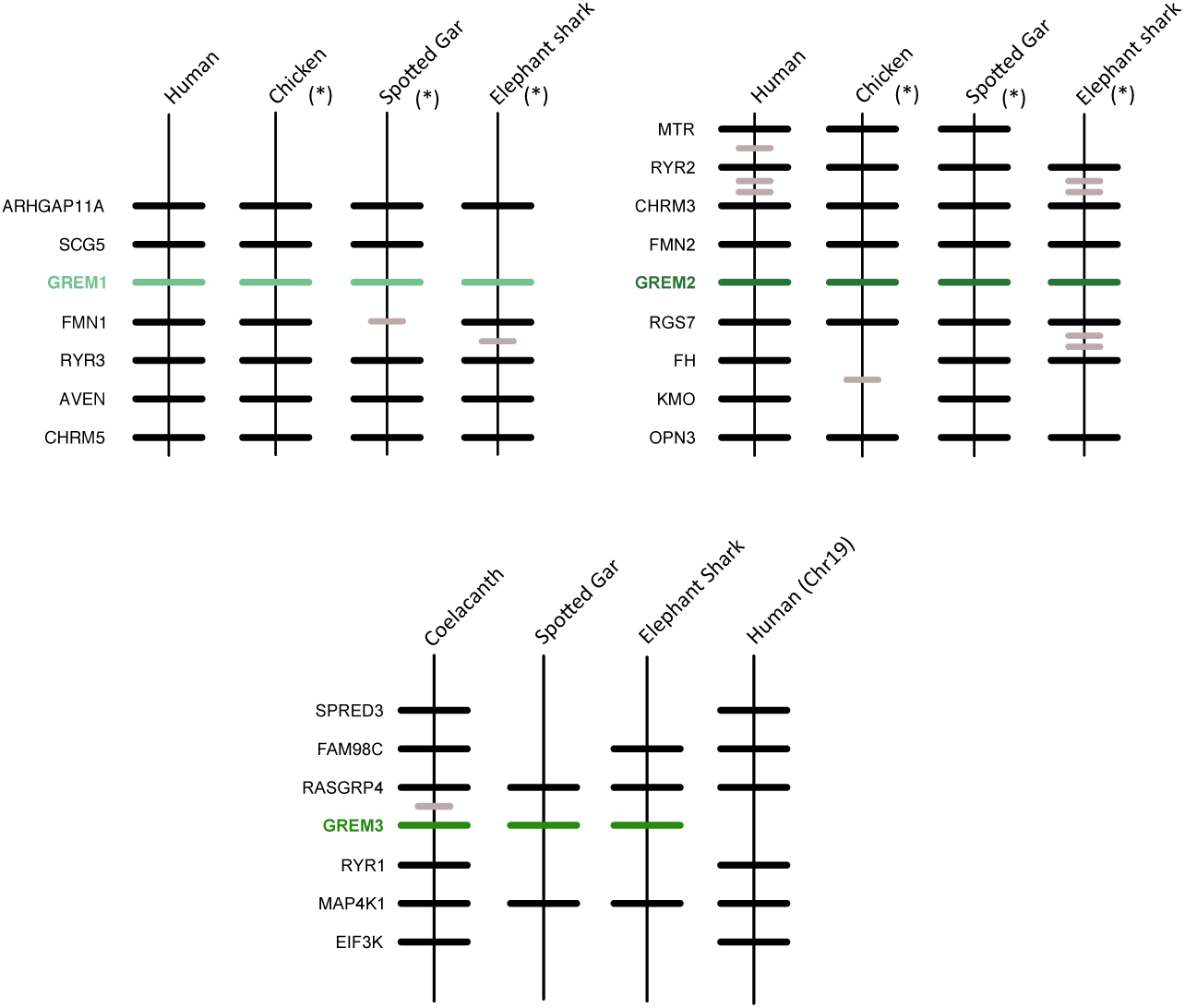
Patterns of conserved synteny in the chromosomal regions that harbor GREM genes. Asterisks denote genomic segments oriented from 3′ to 5′, gray lines represent intervening genes not contributing to conserved synteny, whereas the lack of a line denotes the absence of the gene.

It is necessary to highlight that although we report disagreement in tree topologies between published studies and our results, phylogenetic trees are not always directly comparable as they have differences in the taxonomic and/or membership sampling, two factors known to affect phylogenetic inference.

### Definition of ancestral gene repertoires

The interpretation of the phylogenetic distribution of genes in combination with the evolutionary relationships among them (i.e. gene tree), in the light of the organismal phylogeny (i.e. species tree), represents an appropriate way to understand the evolution of the DAN gene family. According to our results, GREM1, GREM2, SOST, SOSTDC1, and NBL1 are present in the major groups of gnathostomes (Fig. 4; Supplementary Fig. S1E), and within each paralog clade the species arrangement does not significantly deviate from the organismal phylogeny. This indicates that all these genes were present in the common ancestor of the group, and agrees with the expectations of the simplest model of multigene family diversification, the divergent evolution model (Nei et al. 1997). In the case of GREM3, CER1, and DAND5, although they are not found in all major groups of gnathostomes, their phyletic distribution in combination with the corresponding tree topology (Fig. 4) indicate that these genes were also present in the common ancestor of gnathostomes. Thus, our results suggest that the last common ancestor of gnathostomes, dated between 615 to 476 mya (Hedges et al. 2015), possessed a repertoire of eight DAN genes: GREM1, GREM2, SOST, SOSTDC1, NBL1, GREM3, CER1, and DAND5. We note that the ancestral condition of eight genes is only present today in coelacanths (Fig. 4). This reconstruction holds under any of the alternative rootings previously proposed in the literature.

**Figure 4.**
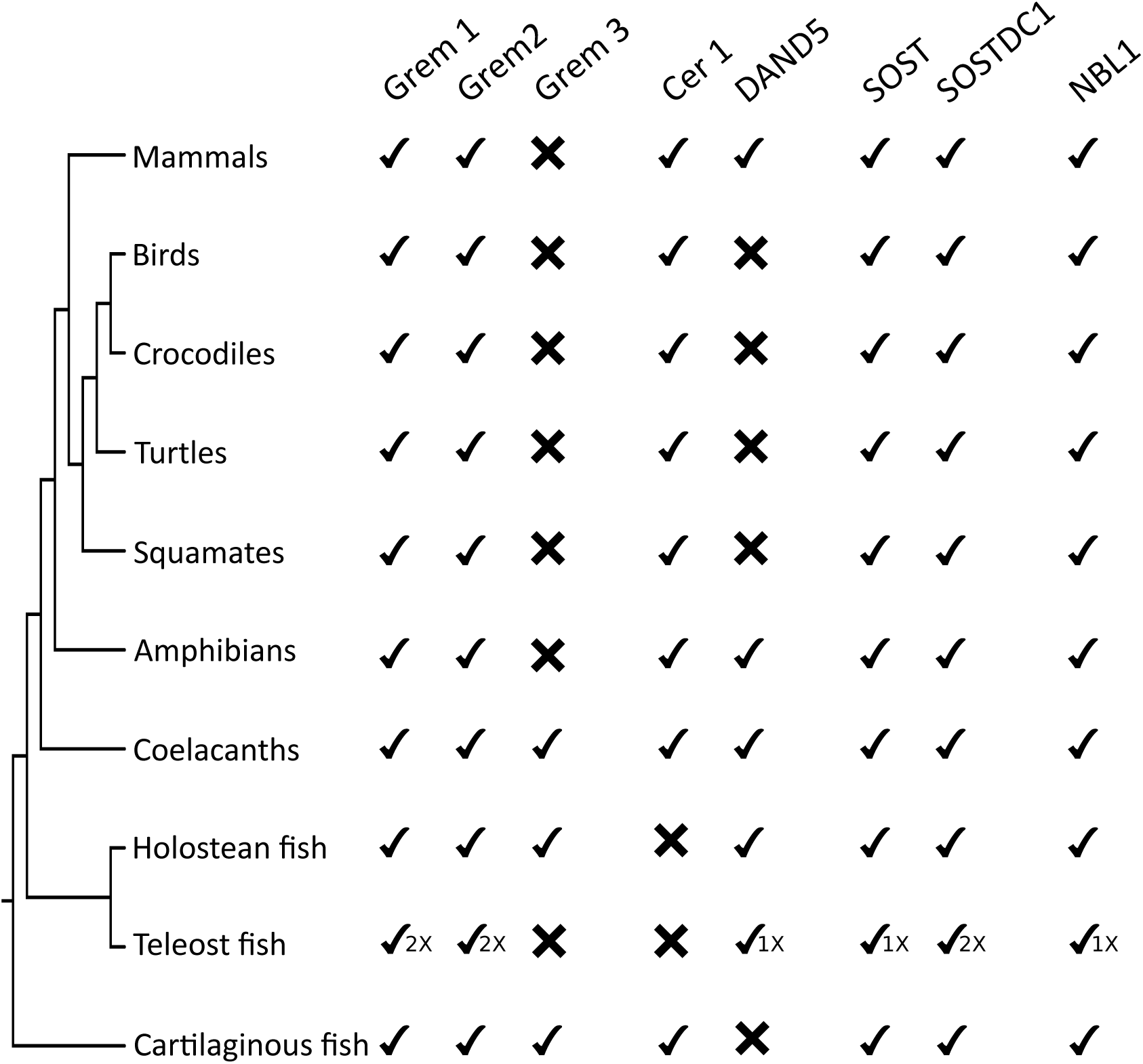
Phylogenetic distribution of DAN genes in gnathostomes.

Inferences about the repertoire of the common ancestors of vertebrates, olfactores and chordates are somewhat speculative given the limited availability of genomes and the quality of current assemblies. However, the phylogenetic position of key taxonomic groups, such as cyclostomes, urochordates, and cephalochordates, provides significant information to advance hypotheses regarding the complement of genes in these different ancestors. We identified GREM2 and SOSTDC1 sequences in cyclostomes (Supplementary Figs. S1A and S1D), indicating that at least these two genes were likely present in the common ancestor of vertebrates. However, given that cyclostome sequences are nested within the GREM2 and SOSTDC1 clades and not sister to a clade with more than one DAN gene, it is possible that the absence of the other paralogs in cyclostome genomes is due to gene loss. Inferences about the common ancestor of olfactores are restricted given the limited number of urochordate genomes and the quality of their assemblies. To further complicate this situation, urochordates have experienced an extraordinary number of gene losses (Dehal et al. 2002; Cañestro et al. 2003; Cañestro et al. 2013; Albalat & Cañestro, 2016), so it is difficult to tell whether the absence of a gene is due to the quality of the data or a real loss. Taking all these uncertainties into account, we were able to identify one urochordate sequence. This sequence was recovered as sister to three cephalochordate GREM genes, which in turn was recovered as sister to GREM2 (Supplementary Fig. S1D). Given that GREM genes diversified as a product of the two rounds of whole genome duplications in the last common ancestor of vertebrates (Singh et al. 2015; Sacerdot et al., 2018; Simakov et al. 2020), a single annotated gene copy in urochordates is expected. Thus, the most probable scenario is that the common ancestor of olfactores possessed one GREM gene. It is not possible to draw inferences for the other members of the gene family at this time. However, given that the ancestor of olfactores existed before the two vertebrate-specific whole genome duplications, we expect its repertoire to have fewer genes compared to the ancestors of vertebrates or gnathostomes.

Finally, we found eight cephalochordate sequences that help us define the repertoire of genes in the ancestor of chordates. One sequence was recovered sister to a lamprey SOSTDC1 gene, a clade that in turn was placed sister to all other SOSTDC1 sequences (Supplementary Fig. S1A). A group of two sequences was recovered sister to the clade that includes NBL1 and GREM genes, and we think this clade represents an ancient member of the gene family that was only retained in this group. Two additional sequences were recovered as sister to the NBL1 lineage; whereas the last three were placed sister to a urochordate GREM sequence, and this latter clade, in turn, was recovered sister to GREM2 (Supplementary Fig. S1C). Our results agree with Le Petillon et al. (2013), who also found copies of NBL1 and GREM orthologs in cephalochordates. However, in contrast to Le Petillon et al. (2013), we did not find a cephalochordate DAND5 gene lineage but instead identified a cephalochordate-specific gene lineage (Fig. 1, purple clade). Bringing these results together, we propose that the chordate ancestor had at least four DAN genes (SOSTDC1, NBL1, GREM, DAND5 and the cephalochordate-specific gene lineage). Le Petillon et al. (2013) also identified sequences in ambulacraria, the group that includes echinoderms and hemichordates, suggesting that the deuterostome ancestor, which existed between 797 and 684 mya (Hedges et al. 2015), had at least three DAN sequences, NBL1, GREM, and DAND5.

### Differential retention of DAN genes during the evolutionary history of gnathostomes

Our results show that copies of GREM1, GREM2, SOST, SOSTDC1, and NBL1 have been maintained in the genome of all major groups of gnathostomes during the last 615 mya (Hedges et al. 2015; Fig. 4), which could suggest they play critical roles during development. In line with this suggestion, the inactivation or deletion of some of these genes produce embryonically lethal effects (Khokha et al. 2003; Gazzero et al. 2006); in other cases, the loss of a gene is associated with pathological conditions (Li et al. 2008), although there are other examples where knockouts develop minor morphological anomalies with no significant consequences (Davis et al. 2015; Voguel et al. 2015). Although these genes are present in all major groups of gnathostomes, some of them could still be absent in a particular group within these more inclusive taxonomic categories. For example, it has been suggested that the GREM2 gene was lost in the common ancestor of ruminants, hippopotamuses, and cetaceans between 56.3 and 63.5 million years ago as a product of a chromosomal rearrangement (Opazo et al. 2017).

Our results also revealed that GREM3, CER1 and DAND5 were differentially retained during the evolutionary history of gnathostomes. The differential retention of genes could be a stochastic process, in which the resulting differences in gene complement do not translate into functional consequences. This phenomenon could be facilitated by a degree of redundancy that could function as a backup with functionally overlapping paralogues, in case that one of the genes is lost or inactivated (Gitelman, 2007; Cañestro et al., 2009; Félix & Barkoulas, 2015; Albalat & Cañestro, 2016). However, it is also possible that possessing multiple copies could help direct the trajectory of physiological evolution by providing opportunities for the emergence of biological novelty (Ohno et al., 1968; Ohno, 1970; Force et al., 1999; Hughes, 1994; Zhang, 2003). For example, the differential retention and duplication of the γ-globin genes might have played an important role in the evolution of life history in anthropoid primates by facilitating an extended fetal development (Goodman et al. 1987; Opazo et al. 2008). Similarly, the differential retention of functional copies of the INSL4 gene may have been associated with unique reproductive characteristics in catarrhine primates (Arroyo et al. 2012a,b).

From another perspective, the absence of genes in natural knockouts represents an opportunity to understand their physiological role (Albertson et a. 2009). This suggestion is based on the orthology-function conjecture: the expectation that orthologous genes are most likely to have equivalent functions in different organisms (Altenhoff et al. 2012). This approach has been useful for understanding human diseases, especially for those exhibiting simple Mendelian inherence, such as cystic fibrosis, albinism and other diseases (Albertson et al. 2009). For example, it has been shown that in the blind cavefish (*Astyanax mexicanus*), loss of pigmentation is produced by mutations inactivating the protein encoded by the OCA2 gene, which is also the most frequently mutated gene in cases of human albinism (Protas et al. 2006). Thus, both retention and gene loss could be seen as evolutionary events that help us to understand the genetic bases of biological diversity.

Our results point to the GREM3 gene as the most extreme case of differential retention among DAN genes. It has been retained in cartilaginous fish, holostean fish and coelacanths (Fig. 4), suggesting that it was present in the common ancestor of gnathostomes, between 615 and 473 mya (Hedges et al. 2015), and was subsequently lost independently in the common ancestors of tetrapods and teleost fish (Fig. 5). A similar case of differential retention of a newly discovered gene lineage was shown in a tumor suppressor gene family (Wichmann et al. 2016). In this case, in addition to the elephant shark, spotted gar and coelacanth, the newly discovered gene lineage was also retained by teleost fish (Wichmann et al. 2016). The absence of GREM3 in a significant fraction of gnathostome species suggests that GREM3 could be dispensable. However, given that GREM3 represents a new gene lineage of unknown biological function, it is premature to evaluate the physiological and evolutionary impacts of retaining or losing this gene. Experimental evidence for the other members of the clade (GREM1 and GREM2) provides conflicting results (Khokha et al. 2003; Davis et al. 2015; Voguel et al. 2015), making it difficult to anticipate the degree of dispensability of the related GREM3 gene.

**Figure 5.**
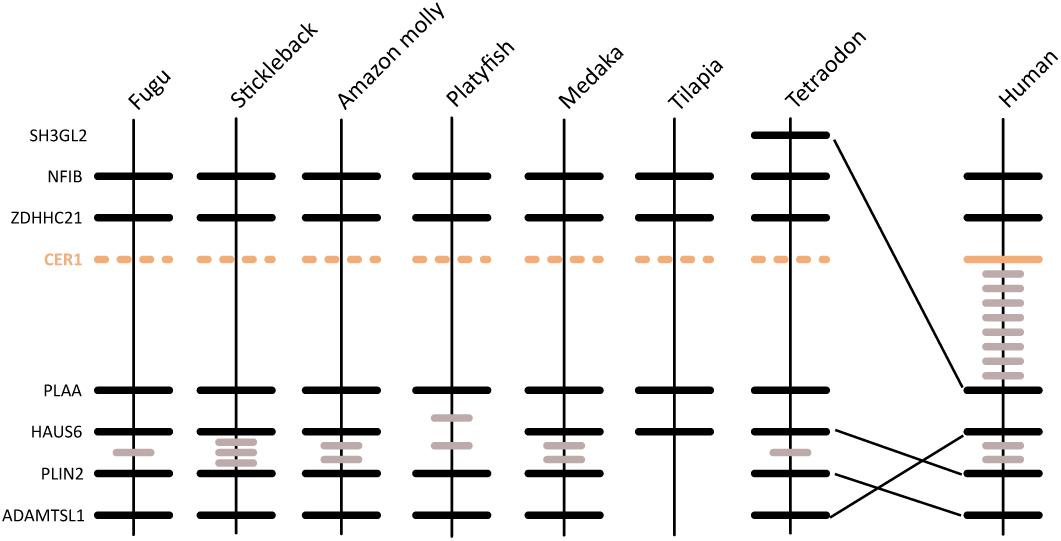
Patterns of conserved synteny in the putative genomic location of the CER1 gene in ray-finned fish. Gray lines represent intervening genes not contributing to conserved synteny, the lack of a line denote the absence of the gene, whereas segmented lines denote the putative location of a missing gene.

CER1 was also present in the common ancestor of gnathostomes and was differentially retained during the radiation of the group (Fig. 4), being lost in the common ancestor of ray-finned fish (Actinopterygii) between 425 and 375 mya (Betancur-R et al. 2013)(Fig. 4). Synteny analyses provide further support for our scenario because the genomic context of CER1 is conserved in fish (Fig. 5). CER1 is a protein possessing nine conserved cysteines and a cysteine knot region expressed during early gastrulation in the primitive endoderm and subsequently in somites and anterior presomitic mesoderm (Shawlot et al. 1998; Pearce et al. 1999; Piccolo et al. 1999; Belo et al. 2000). CER1 has developmental roles associated primarily with anterior-posterior, left-right, and dorso-ventral asymmetries (Belo et al. 2009), accomplishing them by binding to BMPs, Nodal and Wnt ligands, which in turn block the ligand-receptor interaction and activation (Bouwmeester et al. 1996; Piccolo et al. 1999; Belo et al. 2000; Silva et al. 2003; Avsian 2004). The physiological significance of this gene seems to vary with the taxonomic group: for example, CER1 produces stronger phenotypic effects in amphibians than in mammals (Simpson et al. 1999; Piccolo et al. 1999; Belo et al. 2000). In fish, CER1 developmental functions may be performed by its paralog DAND5, which is functionally similar (Hashimoto et al. 2004; Marques et al. 2004; Belo et al. 2009; Lopes et al. 2010; Schweickert et al. 2010; Hamada et al. 2012; Araujo et al. 2014). Thus, from an evolutionary perspective, the possession of both paralogs, CER1 and DAND5, in gnathostomes other than fish would represent a case of functional redundancy. Under this scenario, the loss of one of these genes could be compensated by the retention of the other.

In the case of DAND5, we found this gene in all main lineages of gnathostomes other than sauropsids and cartilaginous fish (Fig. 4). In the case of the elephant shark, we believe that the absence of the gene could be due to a problem in the current genome assembly; accordingly, our proposed scenario mainly assumes the absence of this gene in sauropsids. Our results indicate that DAND5 was lost in the common ancestor of sauropsids, the group including birds and non-avian reptiles, between 312 and 280 mya (Hedges et al. 2015; Fig. 5), even though the corresponding genomic region is well conserved (Fig. 6). Our proposed loss of DAND5 in birds has also been noticed by other authors (Le Petillon et al. 2013). DAND5 has been described as a gene playing fundamental roles driving asymmetries in the early stages of development by inhibiting Nodal activity (Marques et al. 2004). Although in amphibians, fish, and mammals the expression pattern seems to be different, the molecular mechanism, restricting nodal signaling in the left side of the embryo is conserved (Belo et al. 2009). In addition to the roles in early development, during adulthood DAND5 also helps promote cancer metastasis, as well as reactivation of metastatic cells in lungs by reversing the ability of BMP to inhibit cancer stem cell function (Gu et al. 2012). Experiments in other organs, such as bone and brain, show that cancer-related functions of DAND5 are specific to the lung (Gu et al. 2012). Consequently, it has been shown that patients expressing high levels of DAND5 have overall reduced survival rates (Gu et al. 2012). The lack of this gene in sauropsids is difficult to interpret; we speculate that, here, the loss of DAND5 is compensated by CER1, removing any physiological consequences and consistent with the idea that gene families possess some degree of redundancy (Gitelman, 2007; Cañestro et al., 2009; Félix & Barkoulas, 2015; Albalat & Cañestro, 2016). Thus, CER1 and DAND5 apparently perform similar physiological roles during development (Hashimoto et al. 2004; Belo et al. 2009).

**Figure 6.**
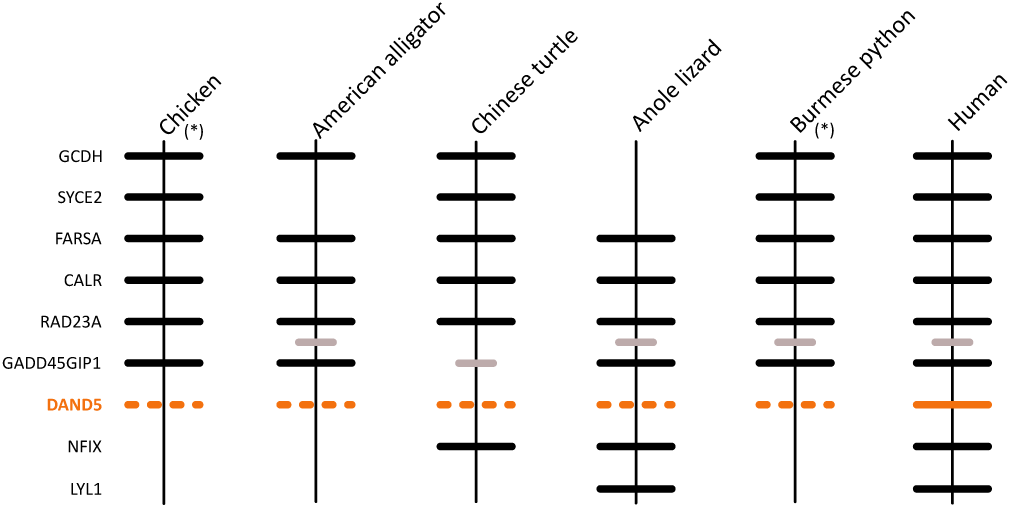
Patterns of conserved synteny in the putative genomic region location of the DAND5 gene in sauropsids. Asterisks denote genomic segments oriented from 3′ to 5′, gray lines represent intervening genes not contributing to conserved synteny, the lack of a line denotes the absence of the gene, whereas segmented lines denote the putative location of a missing gene.

### Rates of molecular evolution: the mammalian slowdown

The phylogenetic trees indicate that the rates of molecular evolution, as measured by branch lengths, vary within each paralog clade (Fig. 1). The most notorious case is the observed slowdown in branches leading to mammals relative to sauropsids (Supplementary Fig. S1A to S1D). This is interesting because sauropsids such as birds, crocodilians, lizards, snakes and turtles tend to have genome-wide rates of evolution much lower than in mammals (Green et al. 2014). Using the relative rate test in MEGA 7 (Kumar et al. 2016) we confirmed that rates in amino acid change are significantly slower in mammals relative to sauropsids (Supplementary Table S2). In mammals, the divergence values ranged from 0.0112 (SOSTDC1) to 0.5143 (DAND5) substitutions per site, whereas in sauropsids they ranged from 0.0326 (GREM2) to 0.9248 (DAND5). In all cases, except for GREM2, the saurospid divergences were higher than in mammals, and the ratio of rates in sauropsids vs. mammals varied from 1.79 (DAND5) to 14.66 (GREM1). We expect genes expressed in early stages of life to be subject to more stringent purifying selection than those expressed in later development, and therefore to have lower evolutionary rates (Goodman, 1961, 1963; Roux & Robinson-Rechavi, 2008). Thus, the slower rate of evolution in mammals could reflect an increased level of functional or structural constraints on these genes. A slowdown in a specific set of genes should be distinguished from a lineage-specific slowdown. In the first case, the reduced evolutionary rate is probably due to differences in the strength of purifying selection between the different lineages, whereas lineage-specific slowdowns are more plausibly be explained by changes in the substitution rate, which could be reduced by lowering of the mutation rate by improved systems of DNA repair mechanisms, longer generation times, or other factors (Goodman, 1985).

## Conclusions

We performed an evolutionary study of the differential screening-selected gene aberrant in neuroblastoma (DAN) gene family in vertebrates. According to our results, this gene family has evolved by a combination of models including divergent evolution, birth-and-death, and differential retention. We recovered the monophyly of all recognized gene family members, which in turn were arranged into five main clades. Importantly, in this work we described the presence of two new DAN gene lineages; one that is only present in cephalochordates (e.g. amphioxus), and other (GREM3) that was only identified in cartilaginous fish, holostean fish and coelacanths. The ancestor of gnathostome vertebrates possessed a repertoire of eight DAN genes. During the radiation of the group, some of them (GREM1, GREM2, SOST, SOSTDC1 and NBL1) were retained in the genome of all major groups, while others (GREM3, CER1 and DAND5) were differentially retained during the evolutionary history of the group. The rate of molecular evolution of mammals is in general low, suggesting an increased evolutionary constraint regime. Finally, we expect that with all these new information researchers in the biomedical field could put their results in an evolutionary perspective.

## Data accessibility

Data and supplementary material are available online: https://zenodo.org/record/3911345#.XvpGJZNKjUJ

## Acknowledgements

This work was supported by the Fondo Nacional de Desarrollo Científico y Tecnológico from Chile (FONDECYT 1160627) and Millennium Nucleus of Ion Channels Associated Diseases (MiNICAD), Iniciativa Científica Milenio, Ministry of Economy, Development and Tourism from Chile to JCO, National Science Foundation (EPS-0903787, DBI-1262901 and DEB-1354147) to F.G.H. S.V.E. acknowledges NSF grant DEB 1355343.Version 3 of this preprint has been peer-reviewed and recommended by Peer Community In Evol Biol (https://doi.org/10.24072/pci.evolbiol.100104)

